# Automated Microbial Classification System based on Deep Convolutional Neural Networks using Images from Colony Picker

**DOI:** 10.1101/2023.06.29.547155

**Authors:** Sehyun Park, Jing Wui Yeoh, Ching Thong Choo, Cheng Kai Lim, Viet Linh Dao, Chueh Loo Poh

## Abstract

Colony screening in single and multi-species environments is an essential step for microbiome studies. However, it possesses a high possibility of inaccurately classifying the species of interest and demands a high degree of manpower and time. Thus, automating the classification of microbes is beneficial to minimize the time and inaccuracy in the colony screening/picking step. Here, we developed an automated microbial classification system for five target species, based on deep Convolutional Neural Networks (CNN) using images captured by an automated robotic colony picker. Multiple possible scenarios of colony culture and diverse morphologies of colonies were examined in building the training and test datasets to train and validate the model and performance on real-life implementations. The final model trained using 60,000 training images, with 12,000 images per species and 3-fold cross-validation, achieved a test accuracy of 94.2% and a test loss of 0.154. Upon testing using a deployment dataset of 4,500 images (900 images per species) with different methods of applying cells onto the agar plate, high accuracy of up to 96.6% was obtained. Five evaluation metrics were implemented to evaluate diverse scenarios of the test data to justify the validity of the model in real-life applications. This model forms a foundation for the classification of more species through transfer learning in the future.

## INTRODUCTION

Microbiome research has gained traction in recent years to unravel how the microbes underpin human health or diseases as well as the ecological system. Recent studies primarily focus on developing analytical approaches, mathematical tools, and engineering solutions to understand the microbiome in the host and to alter the microbial compositions^*1*^. In culturing microbial samples, researchers often need to visually screen the microbial colonies on the agar to discriminate the species colony of interest and picked the colony for further analysis. Colony picking has been applied in various fields of studies including clinical microbiology, biofuels, synthetic biology, food research, etc.^*2,3*^

Colony picking can be done manually or automatically using a robotic platform. Existing studies mostly rely on manual colony picking and screening of the plates in advance of picking the target species since there is no automated classifying tool to identify the specific species^*4*^. However, the manual colony picking process can be a tedious, time-consuming, and labor-intensive task especially when handling hundreds of colony plates in the screening process of most applications. Moreover, it necessitates a substantial amount of experience to accurately identify the targeted species. As a result, both colony picking and classification can be prone to inefficiency and uncertainty. To address these limitations, commercially available automated colony pickers such as PIXL serve to automate the colony picking process. Automated picking is guided by captured images that are processed to pinpoint colonies on the agar for picking. However, these image processing algorithms do not have the capability to classify the species of colonies.

There have been several studies on utilizing machine learning for automated colony/species classification based on imaging, in a concerted effort to enhance screening accuracy and reduce time consumption. A review paper by Kotwal et al. has provided a guideline to highlight steps involved in bacterial classification, which comprised image acquisition, image preprocessing, image segmentation, feature extraction, followed by classification of bacteria^*5*^. Huang et al. compared supervised and unsupervised learning of neural networks for classifying 18 bacterial classes (using 4,892 microscopy images for every bacterial class: 50% for training and 50% for testing) and concluded that the supervised neural networks have promising classification characteristics for bacterial colony pre-screening process with an accuracy of 73% in the classification of 18 bacteria^*6*^. Subsequently, several papers focused on comparing different machine learning techniques such as CNN, support vector machine (SVM), and random forest (RF) to determine the best functioning algorithms specifically for bacterial images. Turra et al. introduced CNN based technique for urinary tract infection (UTI) pathogen classification based on hyperspectral bacterial colonies (using 16,642 colonies images for 8 species) and compared with SVM and RF as feature classifiers, in which the CNN technique achieved the highest accuracy of 99.7%^*7*^. Panicker et al. proposed a CNN model to automate the classification of tuberculosis bacilli from microscopic sputum smear images (1,800 training patches), with three layers of convolutional layers and a fully connected layer for image-denoising and CNN for pixel-level classification, resulting in a recall value of 0.97, a precision value of 0.78, and an F1-score of 0.87^*8*^.

Despite several research works focusing on automating the classification of various bacterial species^*5-9*^, the inconsistency in image acquisition tools (e.g., hyperspectral imaging or digital microscopic imaging) and image segmentation techniques make the model difficult to be generalized for other studies or for commercial use. Additionally, the majority of research conducted utilized imbalanced datasets and limited performance evaluation metrics, for instance, only accuracy was used as an sole index for assessing the model validity^*5*^. However, accuracy alone cannot adequately reflect the false positive and false negative, which are critical in addressing the model’s malfunction and potential instability. Furthermore, most studies only focused on building the model rather than considering the entire process and proposing the practical scenarios for applying the classification model.

In this study, we developed an automated microbial classification system for five target species: *Bacillus subtilis, Pseudomonas putida, Lactobacillus plantarum, Escherichia coli, and Saccharomyces cerevisiae*, based on a deep Convolutional Neural Network (CNN) using images captured by the PIXL colony picker working with a robotic arm ROTOR (Singer Instruments, UK). Multiple possible scenarios of colony culture and diverse morphologies of colonies were incorporated in developing the training and testing datasets to prevent model overfitting with improved generalization and to oversee performance on practical implementations. The final model consisting of balanced data with 12,000 images for each individual species (60,000 total images) was used in model training, cross-validation and hyperparameter optimization and 15,000 images were used as test data. The trained model was then deployed to predict colonies (a total of 4500 images with 900 images for each species) grown in various environments including multi-species colonies, different pinning methods, incubation time, and types of agar plate to test the validity and reliability of the classification model in practical colony picking applications. Five evaluation metrics have been implemented to better evaluate the model performance in microbial classification. The results shows that the classification system achieved a high classification performance for the test data (accuracy ≥ 90%; F1-score ≥ 0.9 for all species) and deployment data with higher diversity (accuracy ≥ 80%; F1-score ≥ 0.82 for most species). The presented automated image classification system based on images acquired by commonly used digital camera implemented in colony picker paves the way towards future colony picker integrated with ML powered classification system for intelligent automated colony screening/picking.

## RESULTS

### Overall workflow

Figure 1 illustrates the workflow established for the automated microbial classification process based upon CNN, working with PIXL and ROTOR. Five of the species were grown in YPD liquid medium or LB liquid medium overnight. The cells were diluted into equal number of cells based on the OD measurements. To plate the cells, both high-throughput plating of microbes using ROTOR and manual spreading by L-Spreader were conducted on PlusPlate and 90mm round plate. The microbial growth was carried out at either 30°C or 37°C. The cells were grown for 16 hours or 48 hours overnight for the training dataset. Using the PIXL software and ROTOR machine, the images of grown colonies were captured and segmented into single colony. The images of single colonies were subsequently pre-processed by adding padding to be used as a dataset for machine learning. The total images were first separated into 80% of the training dataset and 20% of the test dataset. The dataset trained by CNN model was further split into 80% of training set and 20% for validation set. Different variations of hyper-parameters were examined. After finding the optimal hyper-parameters, the trained model was deployed to predict an independent deployment dataset consisting of colonies grown under various conditions including multi-species colonies, different pinning methods, incubation time, and type of agar plate to legitimate the model validity in real-world applications.

**Figure 1:**
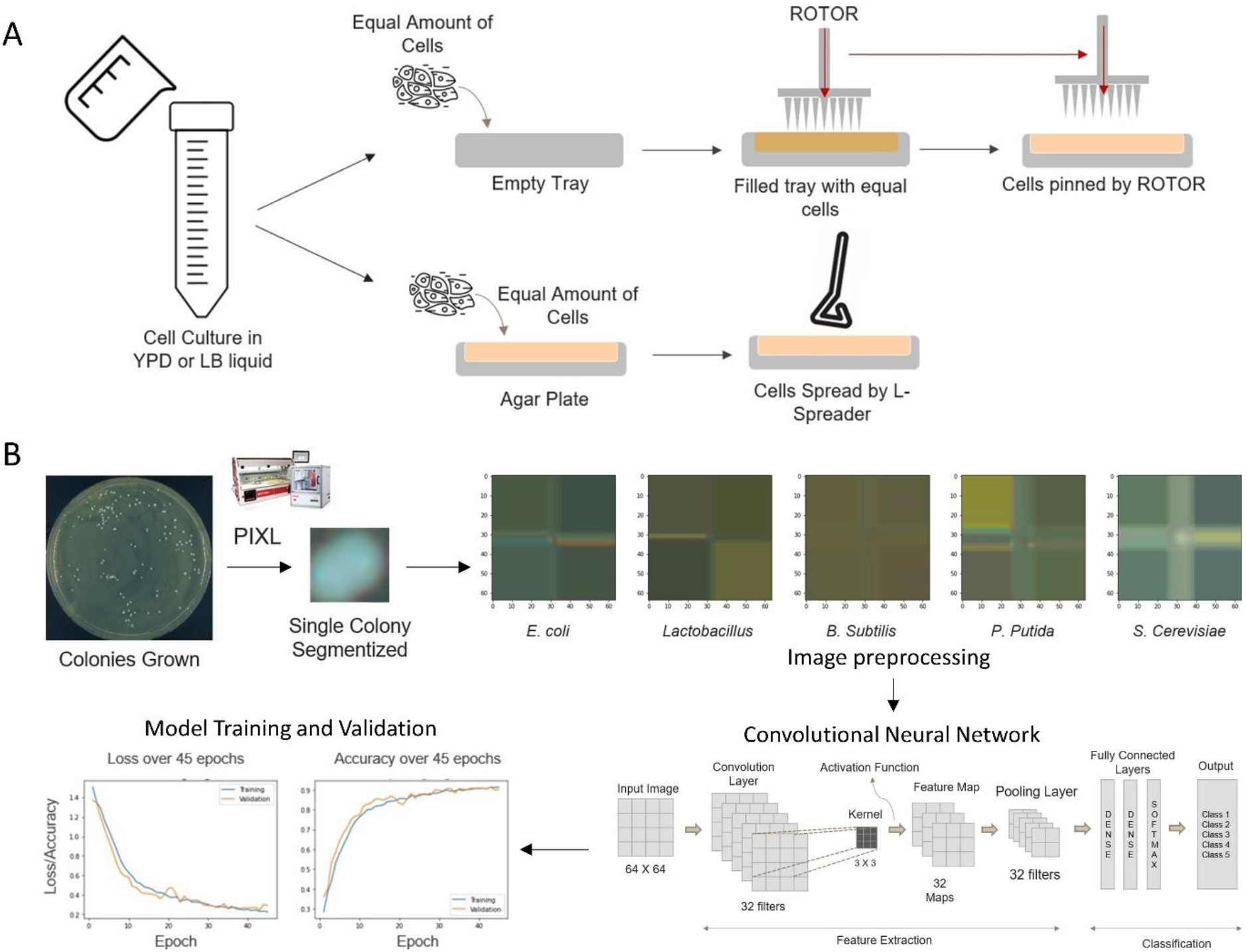
An overview of the workflow. (A) Cell applying methods onto the agar plate: using ROTOR or manually spread using L-spreader (B) Image acquisition and segmentation of colonies using PIXL, followed by image preprocessing by adding paddings for standardization, model training using CNN and model validation.

### Model Training/Development

#### Effect of Increasing training images

The 3-fold repeated cross-validation was implemented for comparing the model performance when training different number of images using the determined optimal hyperparameters. From Figure 2A, with increasing number of training images, we observed an increase in test accuracy and a decrease in test loss. We further examined the precision and recall for the individual species and computed their F1-score. In general, F1-scores for all the species were getting closer to 1 as the number of images increased (Figure 2B). Higher F1-scores implies that the model gives better predictions on classifying the actual label of microbial images with less false positives or false negatives.

**Figure 2:**
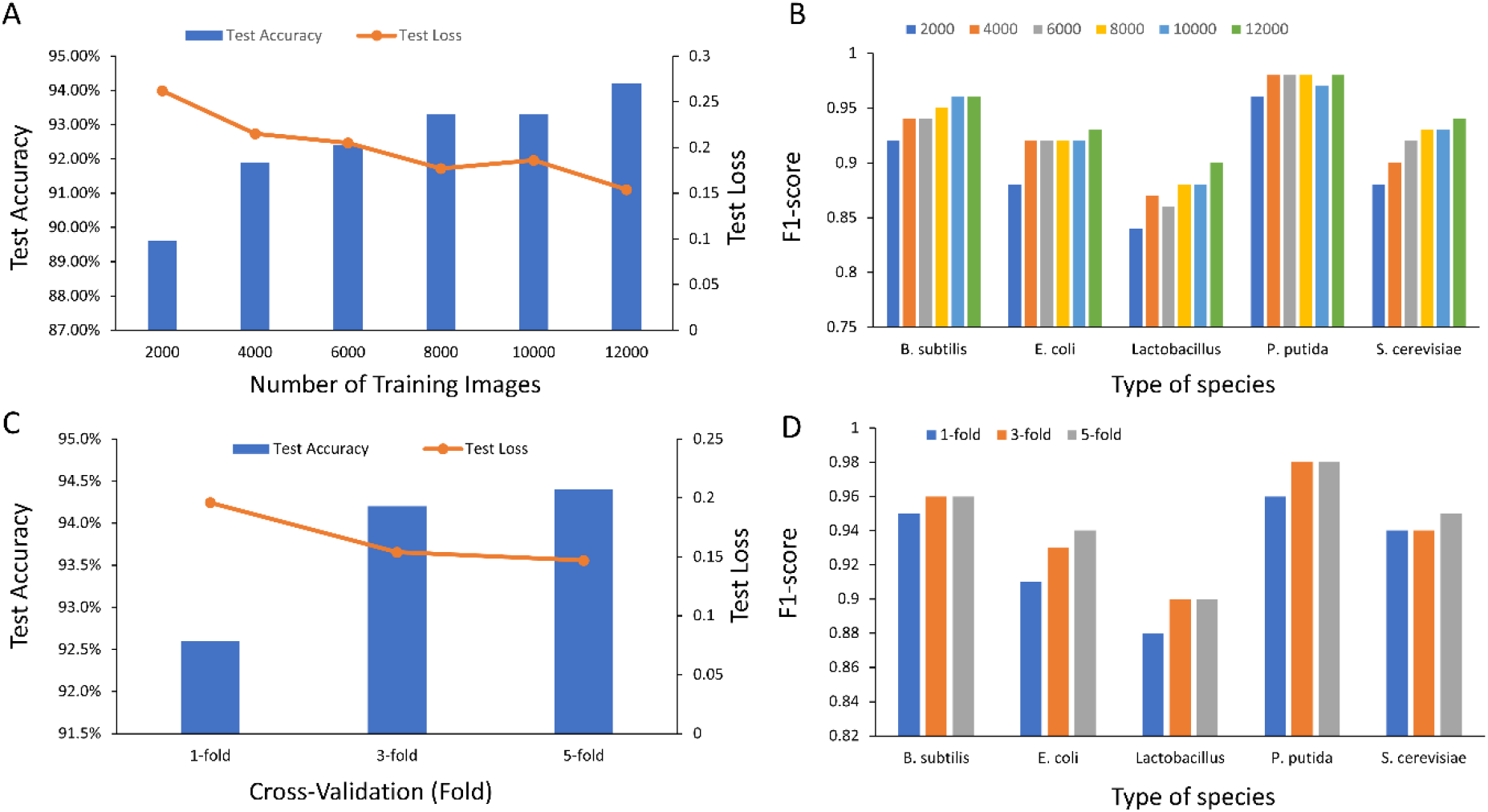
Model performance in response to increasing number of training images and number of folds for repeated cross-validation. (A) The test accuracy (bar) and test loss (line) with increasing number of training images. (B) The computed F1-score based on precision and recall for each species with increasing number of images. (C) The test accuracy and loss with increasing number of folds for cross-validation (D) The computed F1 score across species over increasing number of folds for cross-validation.

#### Effect of folds of cross-validation

For investigating the effect of fold number of repeated cross-validation on model performance, three different k-folds: 1-fold, 3-fold, and 5-fold repeated cross-validation were carried out. As observed from Figure 2C, the test loss and accuracy improved significantly when the number of folds increased from 1-fold to 3-fold. The loss decreased from 0.196 to 0.154 while accuracy increased from 92.6% to 94.2%. However, the test loss as well as test accuracy did not improve considerably when increasing from 3-fold to 5-fold cross-validation. This implies that 3-fold cross-validation could be sufficient to accurately measure the model performance instead of using higher fold cross-validation which would significantly prolong the training duration during hyperparameter tuning. The same phenomena were observed in most of the species when testing the number of folds for repeated cross-validation across different species using F1 score as shown in Figure 2D.

### Model Performance

The trained model was deployed to perform classification on three independent datasets during deployment (not used in training and testing phases): 400 images per species for ROTOR pinned single-species colonies, 400 images per species of manually spread single-species colonies, and 100 images per species of multi-species colonies. For predicting 400 images per species of ROTOR pinned colonies (Figure 3A), the model with 12,000 training images per species outperformed other models, indicating a significant improvement in prediction accuracy to 96.6% with a loss of 0.109. For predicting the type of microbes using 400 images per species of manually spread colonies (Figure 3A), the model with 10,000 training images per species showed the best performance with a prediction accuracy of 90.6% and loss of 0.3794. This is a better result compared to the model with 12,000 training images, which had a loss of 0.486 and accuracy of 88.4%. For the multi-species colonies (Figure 3A), the highest prediction accuracy of 80.0% was achieved when training with 4,000 images per species followed by 10,000 images with 79% accuracy. The lower accuracy could be due to the difference in data distributions since the dataset for the multi-species colonies could be more diverged than the single species training dataset.

**Figure 3:**
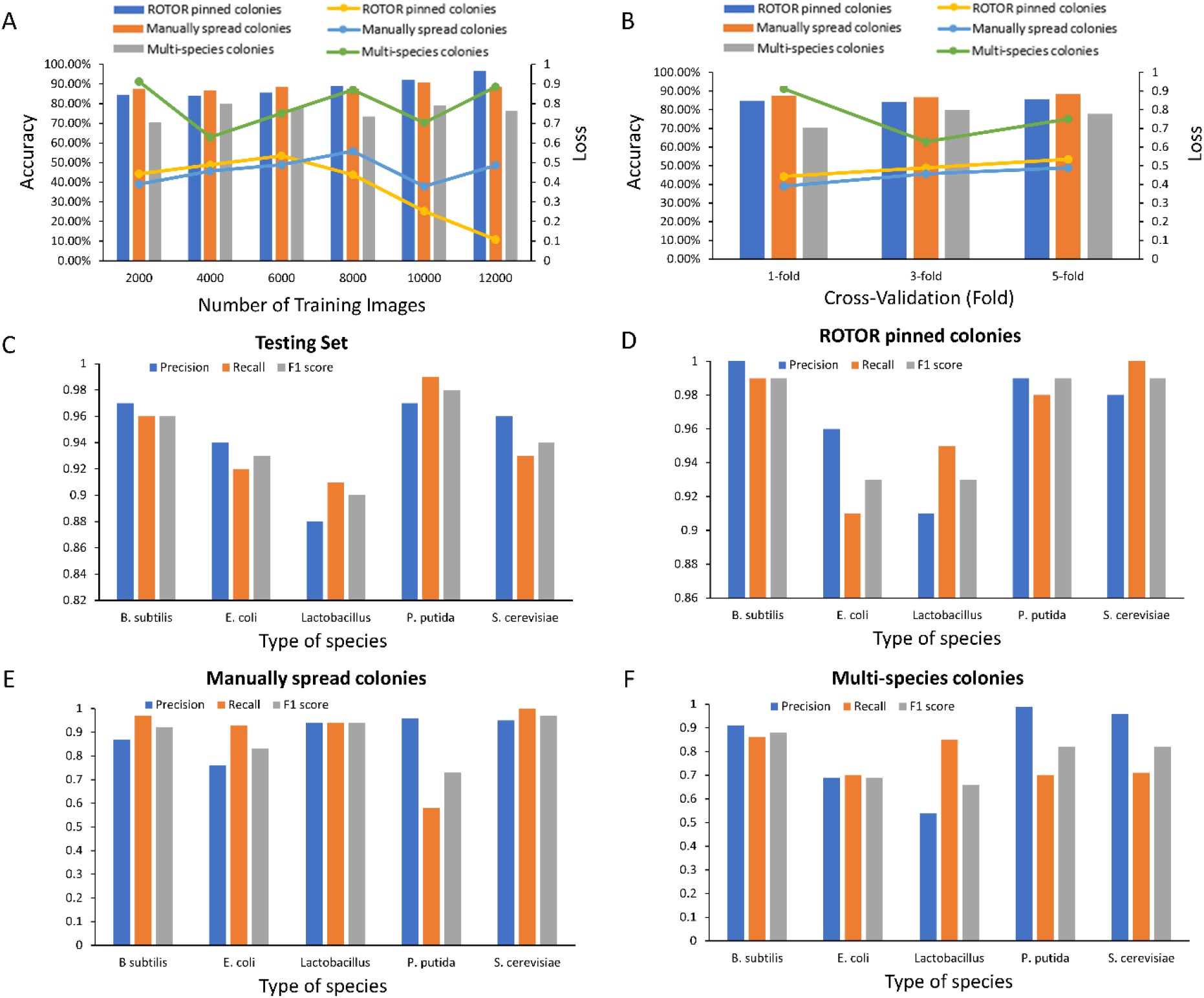
The performance of model prediction on three independent deployment datasets (different methods of applying cells onto the agar plate for single species colonies and multi-species colonies) with increasing number of training images, folds of cross-validation, and across species. (A) Accuracy (bar charts) and loss (lines) for ROTOR pinned single-species colonies, manually spread single-species colonies, and multi-species colonies with increasing number of training images (B) Accuracy (bar charts) and loss (lines) for the three independent deployment datasets over increasing fold of cross-validation. (C-F) Precision, recall, and the computed F1 score with 12,000 training images per species and using 3-fold cross-validation across different species for the (C) testing set (D) 400 images per species of deployment dataset for ROTOR pinned single colonies (E) 400 images per species of deployment dataset for manually spread single colonies (F) 100 images per species for multi-species culture colonies.

The performance of the same three independent datasets used in deployment were assessed with different number of folds of cross-validation. In general, there has been an increase in accuracy with increasing fold, with more significant increase occurring when changing from 1-fold to 3-fold cross-validation for both ROTOR pinned, and manual spread single colonies (Figure 3B). Meanwhile, aside from the accuracy and loss, we sought to assess the other performance metrics: precision, recall and F1 scores of models across the five species with 12,000 training images per species and implemented with 3-fold cross-validation. For the testing set (Figure 3C), all the species exhibited high (≥0.92) values for all the performance metrics except *Lactobacillus* which is in the range of 0.88 to 0.91. While deploying the trained model to predict the performance of 400 images of ROTOR pinned colonies per species (Figure 3D), the performance metrics were considerably higher for the five species (≥0.91) including the *Lactobacillus* (0.91-0.95) though it was at the lowest range of the five species. Intriguingly, when testing with the 400 images per species of manually spread single colonies (Figure 3E), the ranges of performance varied significantly with *E. coli* showed a lower precision value of 0.76 and *P. putida* had a recall value of 0.58, and most of the rest were more than 0.9. For the multi-species colonies with 100 images per species (Figure 3F), the performance is generally lower than the those of the test datasets. The F1 score performance for *E. coli* and *Lactobacillus* was particularly lower, ranging between 0.66 and0.69, while the remaining three species achieved F1 score in the range of 0.82 to 0.88.

## DISCUSSION

In this study, we have developed an automated ML enabled microbial classification system that can identify five target species with high accuracy using data images collected from PIXL colony picker. Diverse scenarios of colony culture and balanced data for the individual species have been employed for the training and testing datasets to avoid model overfitting and to improve model practicality in real-life scenarios. To further validate the model robustness, the trained model was deployed to predict independent data with colonies grown in various scenarios including different pinning methods (automated ROTOR pinned vs manual spread) and multi-species culture colonies etc. and model performance was evaluated in response to increasing number of training images, number of folds of cross-validation, and across species with different performance metrics.

The performance metrics used in evaluating the classification capability of the model are crucial in identifying ways to guide the improvement of model performance. Aside from assessing the common accuracy and loss, it is also important to evaluate the three additional performance metrics such as precision, recall, and F1 score particularly for classification purposes. Precision is measured by counting the number of true positives over the summation of true positives and false positives whereas recall is measured by counting the number of true positives over summation of true positives and false negatives. Low recall values are interpreted as a high number of false negatives or low number of true positives, which are both critical in building a strong predictive model for the microbial classification. This means the model fails to correctly predict the class of the specific species. For example, the low recall value (0.58) of *P. putida* in predicting manually spread colonies (Figure 3E) using model trained with 12,000 training images per species and 3-fold cross-validation is because the model mistakenly predicted *P. putida* as other species rather than correctly recognizing it as *P. putida*. Conversely, the precision value in predicting the same dataset is 0.96, which is significantly higher. High precision with low recall indicates that the number of predictions being classified as *P putida* from the dataset is low while the actual *P putida* colonies are being incorrectly classified as other species. This suggests that the weight and bias gained from the training dataset (precision value of 0.944 and recall value of 0.942 for *P. putida*) is inadequate to capture all the features of *P putida* in the independent dataset during deployment, which could potentially be due to the lack of morphological diversity in the training set relative to other species.

In assessing the performance metrics across the five different species, the precision and recall values of *Lactobacillus* are relatively lower compared to other species in most model evaluations (Figure 3). The precision value of *Lactobacillus* ranges from 0.84 to 0.9, lower than others which tend to be in the range of 0.88-0.98. Also, the range of recall value of *Lactobacillus* is 0.84-0.91, lower than the average range of other species at 0.87-1.00. This may be due to the small size of *Lactobacillus* colonies, resulting in the padding which is used to achieve size of 64×64 dimension dominating the area of processed image.

Analyzing the confusion matrix could derive additional insights into the underlying reasons contributing to most of the observations during the deployment testing. For manually spread single colonies, with increasing fold of cross-validation from 1-fold to 5-fold, as shown in the confusion matrix (Figure 4A), the model learned insignificant features of *P. putida* and incorrectly identified *P. putida* as *E. coli*, resulting in high false negative with low recall for *P. putida*. This phenomenon also emerged when the number of training images increased (Figure 4B). Although correct predictions for *B subtilis, E. coli*, and *S. cerevisiae* increased, wrong predictions for *P. putida* as *E. coli* increased. This is likely because most of the training images of *P. putida* have similar morphologies for ROTOR pinned microbes even in different conditions. Since the training set mostly comprised of ROTOR pinned images, the model might not be able to sufficiently learn the diversified morphologies to classify the *P. putida* that was manually spread by L-Spreader. As the number of training data increased from 10,000 to 12,000 per species, the model might have learned insignificant features that did not correspond well with the features of colonies cultivated from manually spread cells. The false negatives thus lead to moderately lower accuracy at 12,000 images per species.

**Figure 4:**
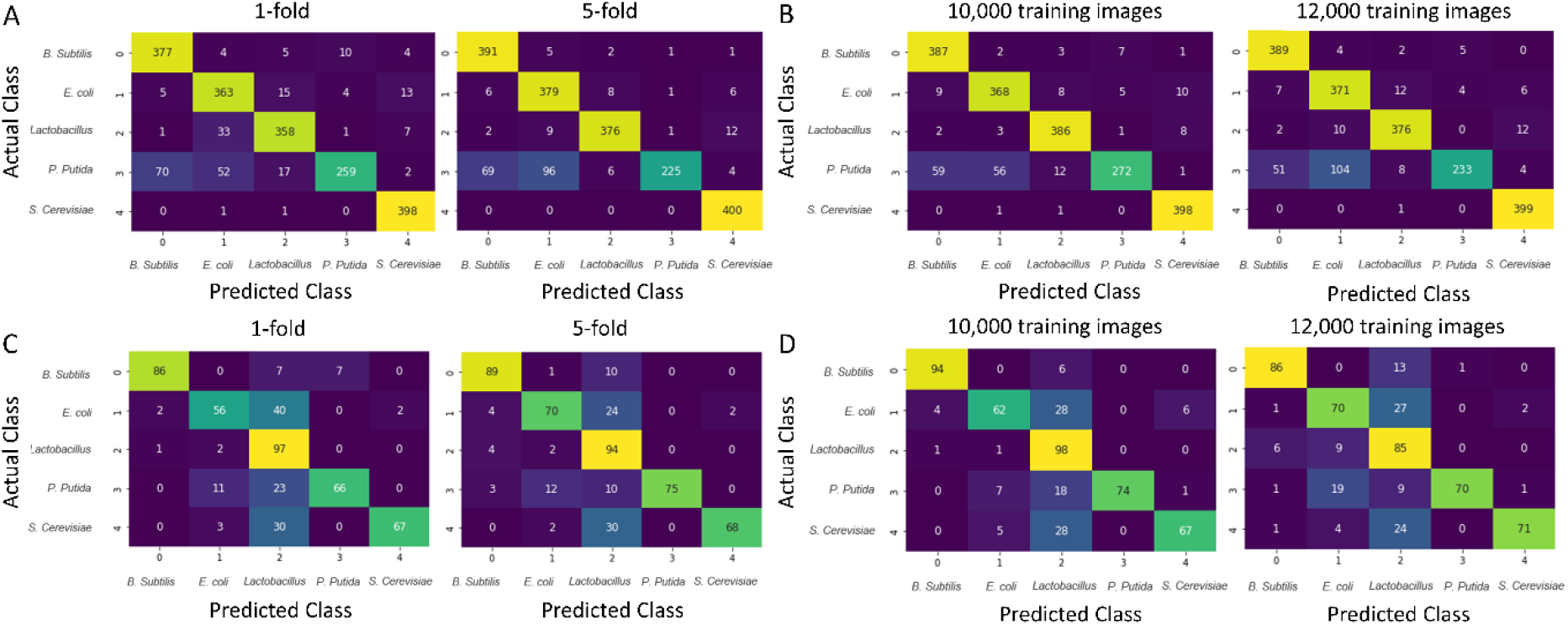
Confusion matrix analysis for the manually spread single-species colonies and multi-species colonies. (A) Comparison of confusion matrix for the five different classes for the manually spread single-species colonies when changing the fold of cross-validation. (B) Performance comparison when increasing the number of training images from 10,000 to 12,000 images. (C) Confusion matrices for the different classes for multi-species colonies when increasing the fold of cross-validation. (D) Performance measurement for multi-species colonies when increasing the number of training images.

Confusion matrix was further utilized to analyze the results for multi-species colonies, particularly the low precision of *Lactobacillus*. While increasing from 1-to 5-fold for the cross-validation, both models exhibited a high number of false positives from other species (Figure 4C). This could be attributed to the tendency of models to predict images with small size colony as *Lactobacillus*. In the multi-species cultures, the different growth requirements, and behaviors of each species of microbe could interfere with the growth of other species, resulting in generally smaller colonies as compared to monocultures. These small colonies were misclassified as *Lactobacillus*, thus increasing the number of false positives of *Lactobacillus*. Meanwhile, the number of true positives decreased when increasing the number of training images from 10,000 to 12,000 for *Lactobacillus, B. subtilis*, and *P. putida* (Figure 4D). This might be due to the variation in colony morphology in multi-species culture and single culture conditions. The second possibility could be the inadequate feature extraction from small-scale images of *Lactobacillus* colonies. As colony images were pre-processed by adding padding, if the image size was too small, there would be more areas with padding. As a result, the extracted features might be repetitive and would be insufficient to determine other important features. The last possibility is that the model might have learned unfavorable features from the images. As the number of images increased by 2,000, there were higher chances of learning insignificant features from the training images which mismatch with features from multi-species culture colonies.

There are multiple challenges when developing and testing the models. One crucial limitation is in the labeling of multi-species colonies. The labelling was largely done by manual classification via visual inspection, which was susceptible to human error. To mitigate this error, reference images of multi-species cultures were assembled and sufficient practices on classifying multi-species colonies were performed prior to the experiment. Meanwhile, it could be enigmatic to compare the results with relevant studies due to the high degree of variations in the dataset, image acquisition method, feature extraction techniques, and classification algorithms between studies. Kotwal et al. provided a comprehensive review on automated bacterial classification using machine learning for the period of 1998-2020^*5*^. The fully automated system using CNN showed good results with an accuracy of 97.3% when trained with bacterial images from Digital Images of Bacteria Species (DIBaS)^*5*^. However, these studies might not reflect the images in real-life implementations with wide diversity. Kotwal et al. also pointed out that the datasets are generally small (<1,000) for deep learning models, and that the models tend to be overfitted or otherwise^*5*^. Unlike the earlier reported studies, the study presented in this paper takes into consideration the various possible scenarios of colony culture and applying diverse morphology of colonies in developing the training, test, and deployment sets to avoid model overfitting and to ensure the model robustness in practical applications.

While the model developed has achieved high accuracy in the automated classification of five microbial species, future work can be done to further improve the model. This could involve enhancing the image resolution, incorporating additional multi-species culture, and introducing different possible culture environments for the training set, which could potentially improve the validation accuracy of multi-species colonies. For augmenting the training images with more variations, a cloud system can be utilized to assemble the colony images captured by PIXL from different laboratories. This could enhance the automation pipeline for colony picking and classification. It is also important to auto-sieve images with unfavorable features to prevent the ML model from learning the noises. Furthermore, the model can be extended to classify more types of microbes. The existing model can serve as a pre-trained model for transfer learning^*23*^, in which new model to classify additional species can benefit from the pre-trained model to reach faster learning on features of new species with high accuracy.

## METHODS

### Colony Culture Environment

All species have their own optimal growth conditions. Understanding the environment for growing colonies of different species is crucial in preparing a dataset with valuable and diverse morphology of colonies. There are numerous factors that could affect the proper growth of microbes, with some of the primary determinants being the incubation time, temperature, and medium composition. There is also a high possibility of imbalanced growth of microbe colonies in limited nutrients^*10*^. Table 1 summarizes the best culture medium, growing temperature, and incubation time recommended by literature. These varying conditions were applied to build the training and testing dataset (Table S1).

**Table 1:** Optimal growing conditions of species

|  | *Escherichia coli*<sup>11</sup> | *Lactobacillus plantarum*<sup>12, 13</sup> | *Bacillus subtilis*<sup>14</sup> | *Pseudomonas putida*<sup>15</sup> | *Saccharomyces cerevisiae*<sup>16, 17</sup> |
| --- | --- | --- | --- | --- | --- |
| Medium | LB broth | MRS, M17 media | LB broth | LB | YPD |
| Temperature (Range) | 37°C (4-45°C) | 35°C (27-34°C) | 30-37°C (18-43°C) | 37 °C (27-41 °C) | 30 °C (20-45 °C) |
| Incubation Time | 8-24 hours | 8-25 hours | 24 hours | 12-72 hours | 5-24 hours |
| MRS (de Man, Rogosa and Sharpe agar)<br>YPD (Yeast Extract-Peptone-Dextrose) |  |  |  |  |  |

### Culture Media

The targeted five species have different growth morphology and optimal environments for colony culture. Referring to literature, various growth environments for culturing microbe species were employed to acquire diverse morphology of colony species. Both YPD and LB were used as cell growth media and for agar plates (Figure S1).

### Spreading Method

To spread the cells onto the agar, three distinct methods were used. Specifically, cells were spread by using L-spreader onto agar on a 90mm round plate, using robotic arm ROTOR to pin the cells to the agar on PlusPlate, or manual pinning of cells to obtain images for various experimental scenarios to be used as training/test or deployment set.

### Cell Dilution

Cells were diluted based on its OD600 measurements to ensure a consistent number of cells were pinned on the agar plate. The details of OD values and sample calculations for each species cultured in YPD and LB liquid are provided in Table S2.

### Image Segmentation and Refinement

The images of grown colonies were captured and segmentized by PIXL with camera resolution of 5MP (2448 × 2048) (Singer Instruments, UK). The PIXL software segments the colonies in different manners based on four algorithms (background subtraction, adaptive detection, colony separation, and specific detection). The adaptive detection algorithm was employed in this study, which showed the best detecting and segmenting capability (Figure S2). To prevent the machine learning model from learning noises and unfavourable features, the images with more than one colony, severe light distraction, large image dimension were removed from the training image dataset. Figure S3 shows the standard colony morphology of the five species and the invalid colony images.

### Dataset

From refined images, the full training dataset of 60,000 images was created by randomly selecting 80% of the data, with 12,000 images per species for the five species. The other 20% of data (15,000 images) were used as the test dataset to assess the model prediction performance. For the independent deployment dataset, microbial images of 4,500 images, with 100 images of multi-species colonies, 400 images of ROTOR pinned colonies, and 400 images of manual spread colonies, for the individual species were used to reflect the real incidences after segmenting images from PIXL.

### Image Preprocessing

The images were processed by adding paddings to standardize the dimension of images to 64×64 while preserving the original size of each colony. From our study, extending and replicating the side pixels as padding showed the best validation accuracy and loss compared to same color padding and no padding (results not shown). Also, we showed that colored images have better validation accuracy compared to grayscale images. As such, extending and replicating the side pixels as padding in color dimension were applied throughout our study (Figure S4).

### Building the Convolution Neural Network (CNN) Model

CNN is a machine learning algorithm that can extract and weigh the important features of the images to differentiate one image from the others^*18*^. It consists of a convolution layer with a convolution filter and the activation function to extract the features from the images, a pooling layer to down sampling the images, and a fully connected layer to take the feature map and generate a probability array indicating the classification of the input images. These layers are stacked into several layers and utilized as both image recognizer and classifier without any handcrafted feature extractor^*19*^. This process is backpropagated each iteration to update weights that indicate the importance of a particular feature in predicting the final output, to minimize the loss function^*20*^. Since the model classifies five species and uses non-one-hot vector labels, sparse categorical cross-entropy was selected as a loss function.

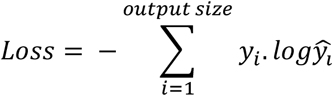

Where *y*_*i*_ is the truth label and *ŷ*_*i*_ is the softmax probability for the i^th^ class.

TensorFlow is an open-source platform for end-to-end machine learning and deep learning tasks, which supports Keras as a high-level neural network library with useful tools for optimization, scoring, and efficient learning of model^*21,22*^. The fundamental structure of CNN model had been created using TensorFlow and Keras. The hyperparameter optimization with cross-validation was then performed to optimize and evaluate the model performance.

### Cross-validation

Scikit-learn is a comprehensive machine learning library in Python that provides an array of tools for machine learning process including data pre-processing, model evaluation, and hyper-parameter optimization. It was used to randomly split the data into 80% of training data and 20% of test data. Next, the training data was further split into 80% of training set and 20% of validation set. In this study, repeated k-Fold cross-validation or repeated random sub-sampling cross-validation was adopted for model evaluation or validation during the training phase. This technique is a variation of standard k-Fold cross-validation in which the k refers to the number of times the model is trained. For every iteration, 20% of samples will be randomly selected as validation set and the rest 80% samples become the training set. The random selection of samples for each iteration makes the repeated k-Fold technique more robust to selection bias. In each fold of the training process, callback tools such as early stopping, and model checkpoint were utilized to inspect how well the trained model predicts the actual label of validation set. Early stopping is a method to stop the training process when there is no further improvement in the model’s validation loss. The model checkpoint saves the best performing model during each epoch in the fold. After all the folds, this model was used to predict the test data and the resulting confusion matrix with true positive, true negative, false positive, and false negative was utilized to better evaluate its performance on the test data.

### Hyperparameter Optimization

Multiple hyperparameters including batch and epoch size, type of optimizers, validation split, number of convolutional layers, size of convolutional layers, dropout rate, size of dense layer, and number of dense layers were tuned and compared to obtain the optimal hyperparameter set (Table 2). The best hyperparameter set is provided in Supplementary Information (Table S3 and Figure S5)

**Table 2:**
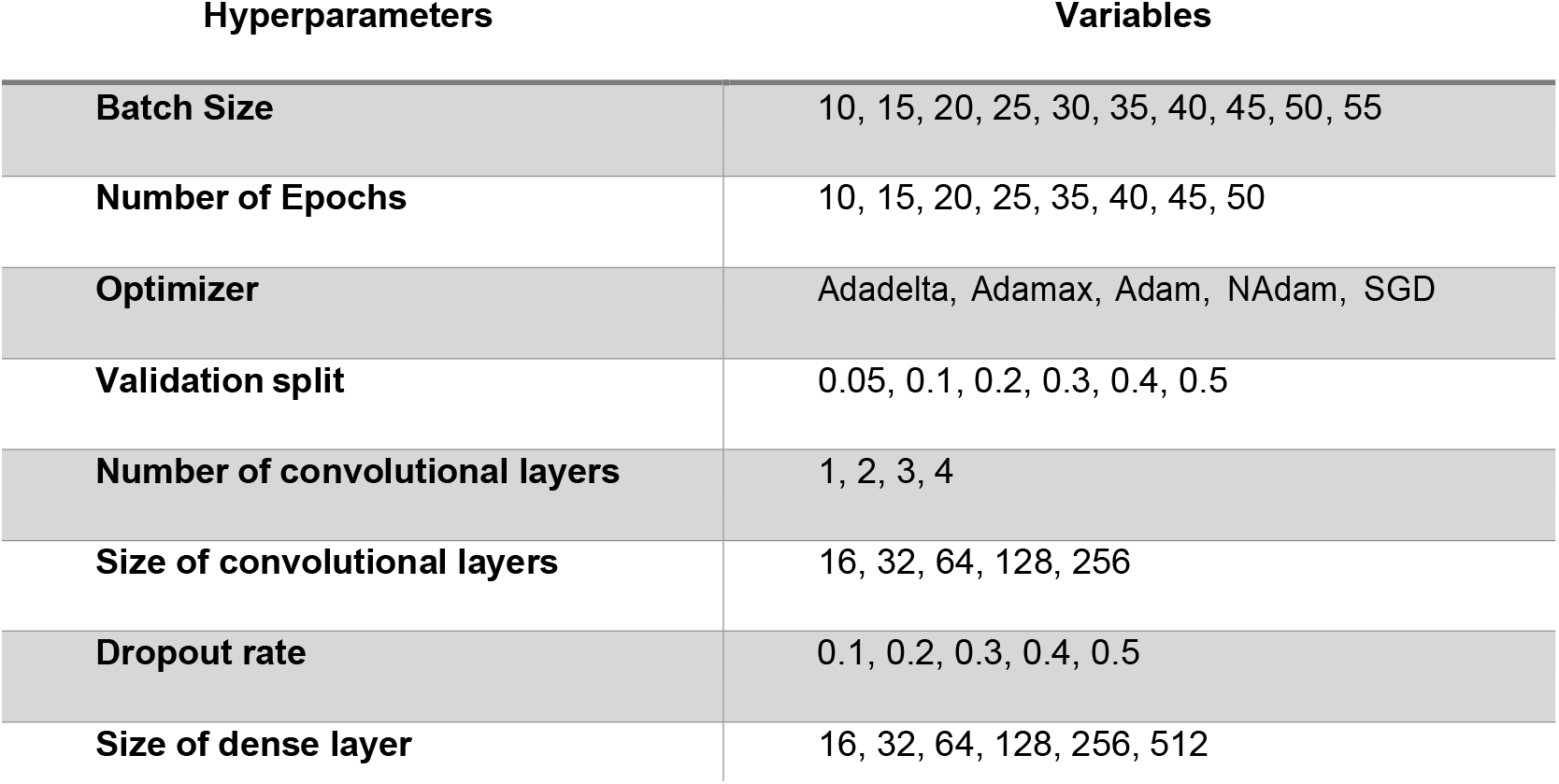
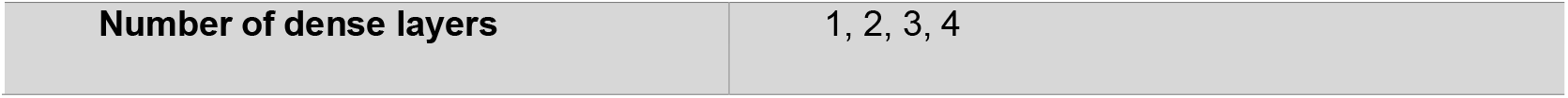
Variables investigated for all the hyperparameters.

| Hyperparameters | Variables |
| --- | --- |
| Batch Size | 10, 15, 20, 25, 30, 35, 40, 45, 50, 55 |
| Number of Epochs | 10, 15, 20, 25, 35, 40, 45, 50 |
| Optimizer | Adadelata, Adamax, Adam, NAdam, SGD |
| Validation split | 0.05, 0.1, 0.2, 0.3, 0.4, 0.5 |
| Number of convolutional layers | 1, 2, 3, 4 |
| Size of convolutional layers | 16, 32, 64, 128, 256 |
| Dropout rate | 0.1, 0.2, 0.3, 0.4, 0.5 |
| Size of dense layer | 16, 32, 64, 128, 256, 512 |
| Number of dense layers | 1, 2, 3, 4 |

### Evaluation Metrics

The model was evaluated based on these indexes: testing loss, testing accuracy, time consumed for complete training of model, and proximity of validation metrics and training metrics. Figure S6 illustrates an example that compared the dropout rates of 0.2 and 0.8. The model performance was also evaluated based on confusion matrix and classification report with evaluation metrics including precision, recall, and F1 score (Figure S7). Confusion matrix compares the actual labels of the images with the predicted labels by the model. In this multi-class classification problem, the matrix contains the number of samples that were correctly classified and the number of samples that were misclassified for the individual classes. Evaluation metrics such as precision, recall, and F1-score can be computed for each class using the values from the corresponding row or column of the confusion matrix. The support refers to the number of images predicted for each individual species.

### Data deployment

Various practical application scenarios were tested to check the validity of the model. For applying cells onto the agar plate, ROTOR pinned single colonies, manually spread single and multi-species colonies were investigated. The conditions used to generate the training set were also implemented to build the deployment set. In addition, the colonies were grown under more diverse conditions, including incubation periods of 8 or 16 hours, multi-species cultures of two distinct species, and on agar plate of different thickness. To this end, the deployment dataset consisted of 4,500 images with each species comprising 100 images of multi-species colonies, 400 images of ROTOR pinned single colonies, and 400 images of manually spread single colonies. The images of colonies grown in multi-species culture were pre-processed by human observer to label the images into correct species after the processing by PIXL (Figure S8).

## CONCLUSION

We have established an automated microbial classification system based on CNN, using 12,000 images per species at diverse conditions captured by PIXL colony picker, to classify five target species: *B. subtilis, P. putida, Lactobacillus, E. coli*, and *S. cerevisiae*. The final model developed using 60,000 training images with optimized hyperparameters and 3-fold cross-validation achieved a test accuracy of 94.2%, loss of 0.154, with an average precision value of 0.944 and average recall value of 0.942 across species during the test phase. The trained model was then deployed to predict colonies grown in various environments including multi-species colonies and different pinning methods at different incubation time, and types of agar plate to justify the validity of model in real applications. From the deployment, the classification system achieved the highest accuracy of 96.6% for ROTOR pinned single culture colonies, 90.6% for manually spread single-species colonies, and 80.0% for multi-species colonies with increasing number of training images. Confusion matrices with three additional performance metrics were further analyzed to gain deeper insights into the performance of individual species, and to identify the potential areas for improvements. This study has presented a non-microscopic image acquisition and classification workflow to automate the colony picking and classification process which could benefit the colony screening process applicable in various fields of studies.

## Author Contribution

S.H.P., C.T.C., C.L.P. conceived the project. S.H.P. and C.T.C. developed the models. S.H.P. and C.T.C. performed the experiments. C.K.L. and V.L.D. advised and guided the experiments. All authors analyzed the results. S.H.P. and J.W.Y. wrote the manuscript with input from C.L.P. All authors commented on and approved the manuscript.

## Acknowledgment

The authors would like to acknowledge the support from the Singapore National Research Foundation [NRF SBP-P5] and the Synthetic Biology Initiative of the National University of Singapore [DPRT/943/09/14]. We would also like to thank Dr Chong Da from Singer Instruments for his advice and assistance on the use of the colony picking instruments and software.

## For Table of Contents Only

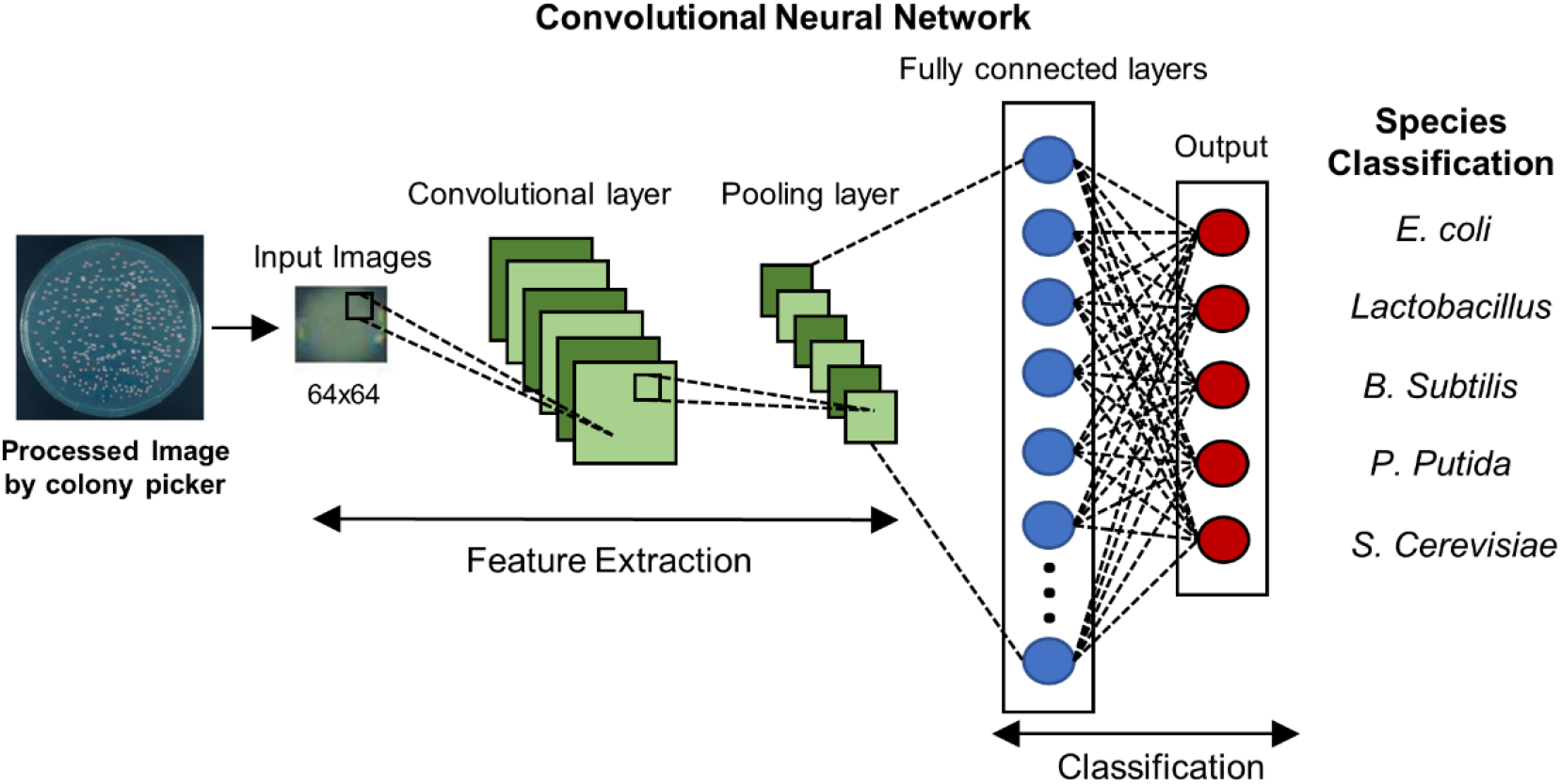

