## Supplementary material for "Automated Microbial Classification System based on Deep Convolutional Neural Networks using Images from Colony Picker": Supplementary_information.pdf

**Table S1:** Variables for colony culture (T: variable for training set; D: variable for test set)

| **Species** | **Plate Type** | **Plate Agar** | **Culture Media** | **Spreading method** | **Cell Culture** | **Cell Dilution Method** | **Incubation Time** |
|---|---|---|---|---|---|---|---|
| *E. coli* | PlusPlate (T, D) | LB Agar | LB Liquid | L-Spreader (T, D) | Single Culture (T, D) | Equal Volume Dilution (x3000, x5000, x1000) (T, D) | Overnight (T, D) |
| *S. cerevisiae* | | YPD Agar | YPD Liquid | Manual Pinning (D) | | | 8hours (T) |
| *Lactobacillus* | 90mm Round Plate (D, T) | M17 Agar (for *Lactobacillus*) | M17 Liquid (for *Lactobacillus*) | Pinning by ROTOR (T, D) | Multi-Culture (D) | Equal Cell number (Based on OD Measurement) (T, D) | 16hours (T) |
| *B. subtilis* | | | | | | | |
| *P. putida* | | | | | | | |

**Table S2:** OD values and sample calculation of each species cultured in YPD liquid and LB liquid.

| | *Escherichia coli* | *Lactobacillus plantarum* | *Bacillus subtilis* | *Pseudomonas putida* | *Saccharomyces cerevisiae* |
|---|---|---|---|---|---|
| OD value cultured in YPD Liquid | 0.1437 | 0.2781 | 0.9874 | 0.1092 | 0.5547 |

| | | | | | |
|---|---|---|---|---|---|
| OD value cultured in LB Liquid | 0.4620 | 0.0241 | 0.6400 | 0.5031 | 0.1694 |
| Number of cells in OD=1 | 2.6x $10^9$ | 6.7x $10^8$ | 1.5x $10^8$ | 3.8x $10^7$ | 3x $10^7$ |

The calculations were based on the following formula:
$(\#\ of\ cells) = (OD\ value) \times (number\ of\ cells\ in\ OD = 1) \times (dilution\ ratio)$

Sample calculation of *E. coli* cells cultured in YPD liquid medium if it is diluted 10x for OD measurement is:

$(\#\ of\ cells) = 0.1437 \times (2.6 \times 10^9) \times 10 = 3.74 \times 10^9\ cells$

*The volume of cell medium can be adjusted by these values.

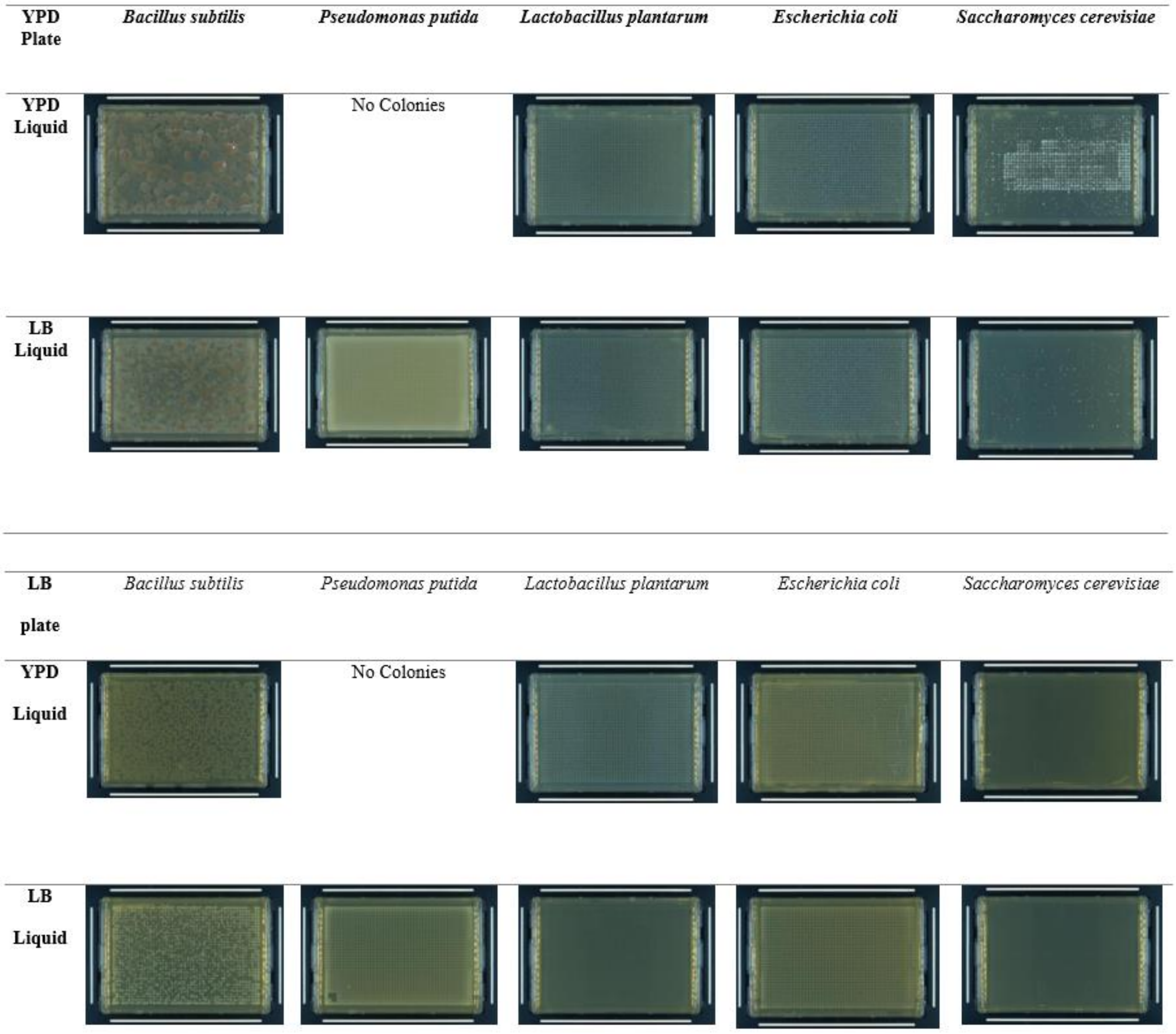


**Figure S1:** Images of colonies cultured under different environments in YPD agar plate and LB plate.

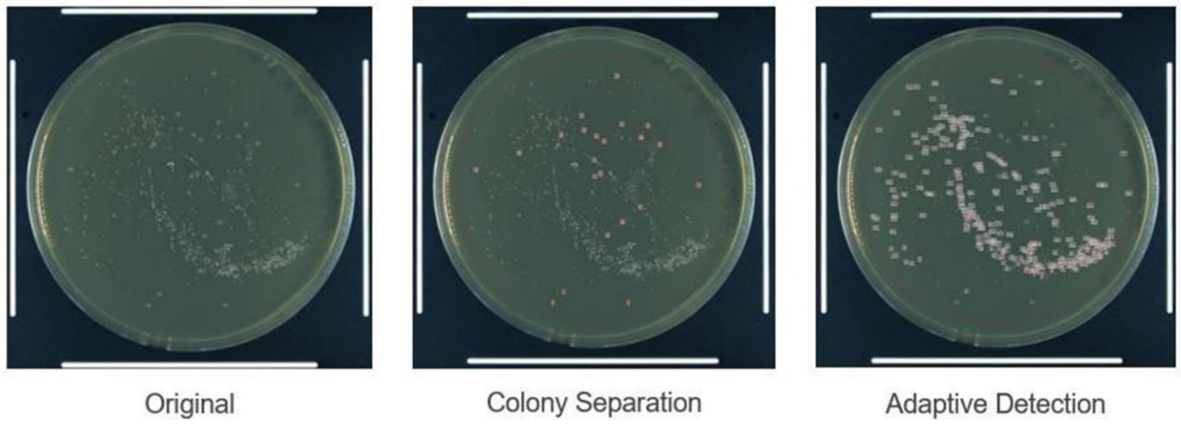


**Figure S2:** The different algorithms of PIXL for colony detection.

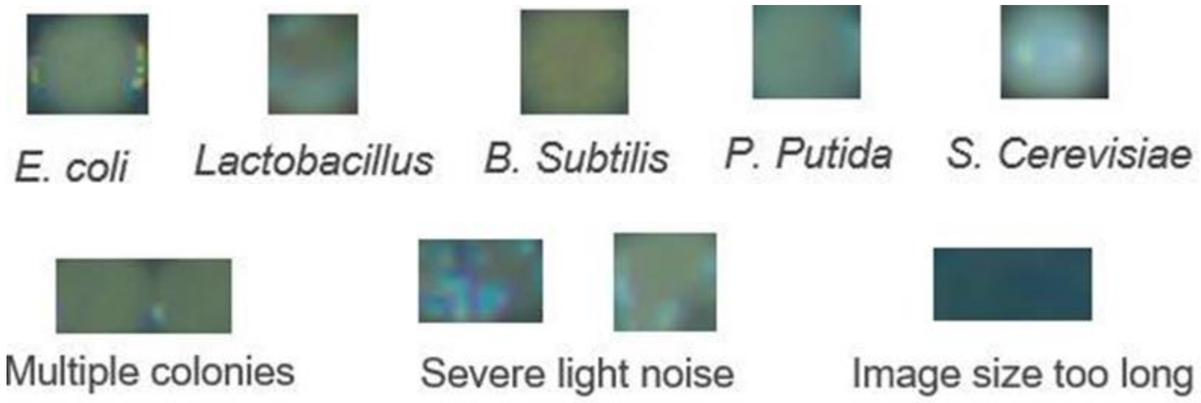


**Figure S3:** Colony morphologies of five species (top panel) and examples of discarded images (bottom panel)

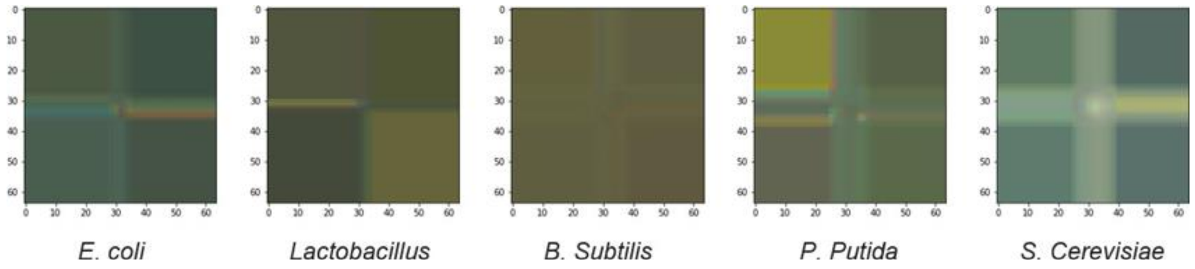


**Figure S4:** Padding added to images for all five species to standardize the dimension of images to 64x64 while preserving the original size of each colony.

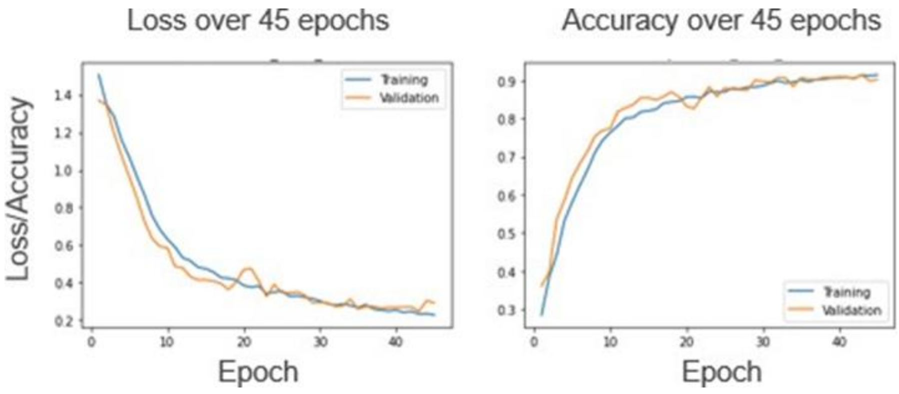


**Figure S5:** Performance metrics (loss and accuracy) after hyperparameter tuning. The optimized hyperparameter was built based upon 1740 images acquired at the start of the project as a training set to minimize the time consumption for calculations. The optimized model shows good proximity of validation metrics and training metrics. The model had training loss of 0.228, training accuracy of 0.914, and time consumed for 45 epochs of 1479.06s.

**Table S3:** Optimized hyperparameter values for the model.

| Hyperparameters | Variables |
|---|---|
| **Batch Size** | 35 |
| **Number of Epochs** | 45 |
| **Optimizer** | Adamax |
| **Validation split** | 0.2 |
| **Number of convolutional layers** | 3 |
| **Size of convolutional layers** | 32 |
| **Dropout rate** | 0.3 |
| **Size of dense layer** | 256-128-64 |
| **Number of dense layers** | 3 |

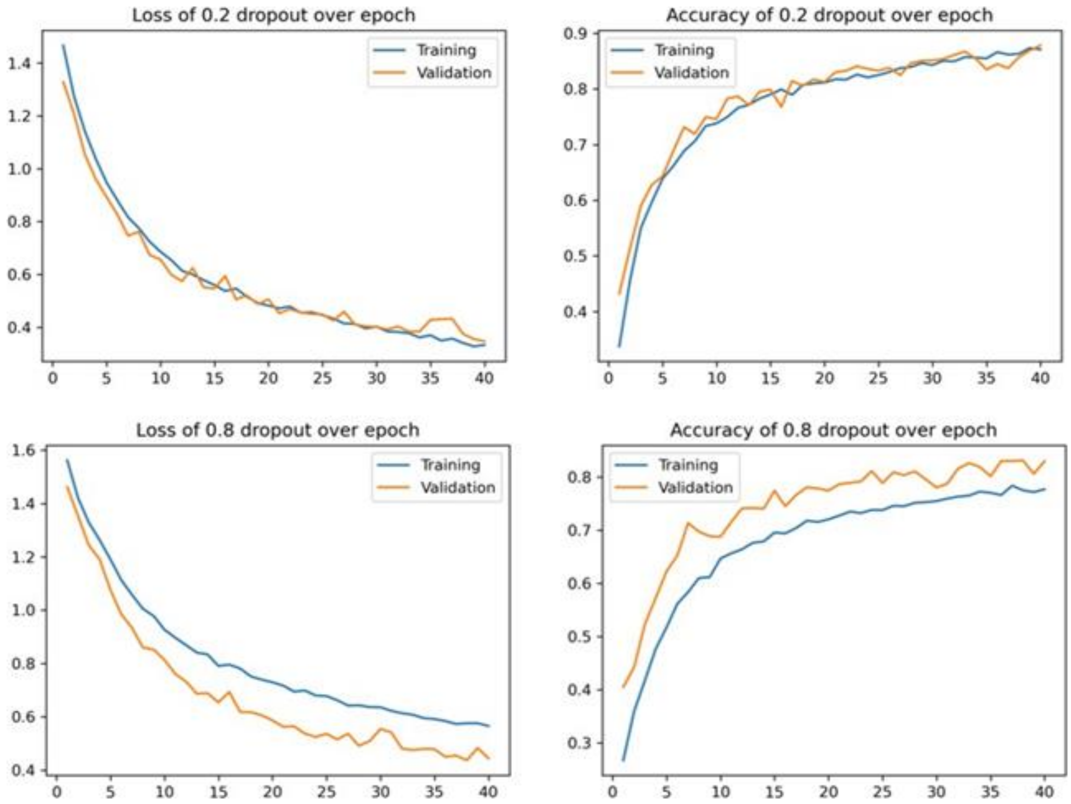


**Figure S6:** The graph showing the performance metrics (loss and accuracy) over epochs for dropout rate (r) of r = 0.2 and r = 0.8. Times consumed for r = 0.2 and 0.8 were 954.78s and 1027.38s respectively.

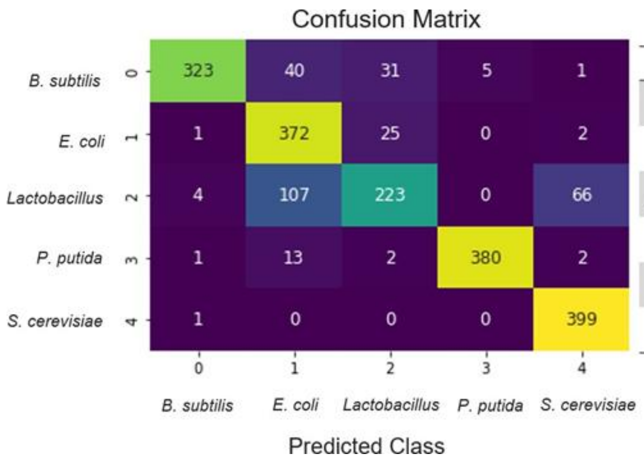


Evaluation Table

| | Precision | Recall | F1-Score | Support |
|---|---|---|---|---|
| ***B. subtilis*** | 0.98 | 0.81 | 0.88 | 400 |
| ***E. coli*** | 0.70 | 0.93 | 0.80 | 400 |
| **Lactobacillus** | 0.79 | 0.56 | 0.65 | 400 |
| ***P. putida*** | 0.99 | 0.95 | 0.97 | 398 |
| ***S. cerevisiae*** | 0.85 | 1.00 | 0.92 | 400 |
| **Accuracy** | | 0.85 | | 1998 |

**Figure S7:** Sample confusion matrix and evaluation table

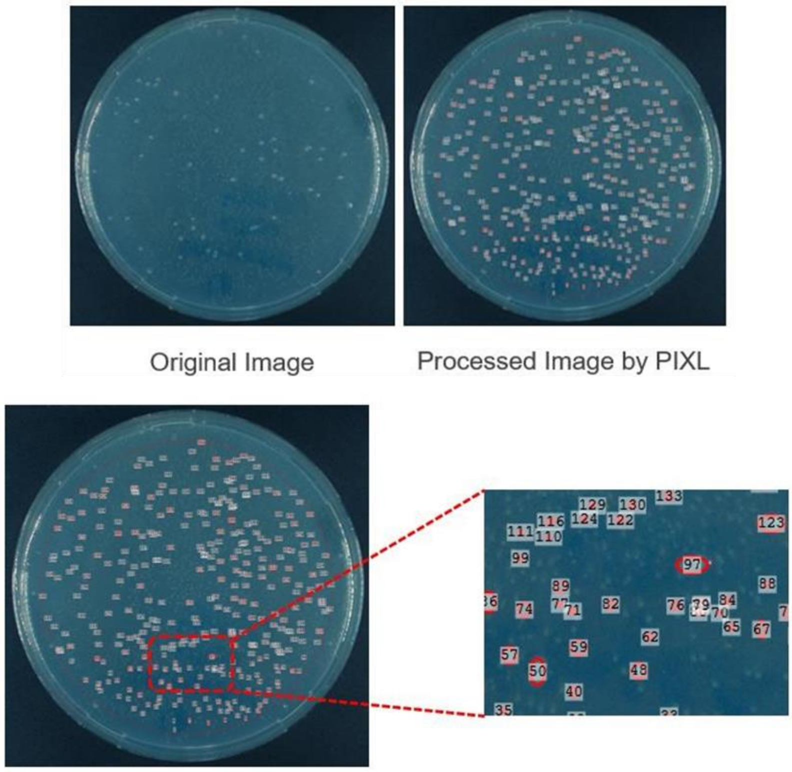


**Figure S8:** Processing of multi-species colonies grown on the YPD agar plate using the algorithm from PIXL.
